# Immune escape mutations selected by neutralizing antibodies in natural HIV-1 infection can alter coreceptor usage repertoire of the transmitted/founder virus

**DOI:** 10.1101/2021.11.10.467978

**Authors:** Manukumar Honnayakanahalli Marichannegowda, Hongshuo Song

## Abstract

The ability of HIV-1 to evade neutralizing antibodies (NAbs) *in vivo* is well demonstrated, but the impact of NAb escape mutations on HIV-1 phenotype other than immune escape itself has rarely been studied. Here, we show that immune escape mutations selected by V3-glycan specific NAbs *in vivo* can alter the coreceptor usage repertoire of the transmitted/founder (T/F) HIV-1. In a participant developed V3-glycan NAb response, naturally selected mutations at the V3 N301 and N332 glycan sites abrogated CCR8 usage while conferred APJ usage on the cognate T/F strain. Mutations at the N301 glycan also impaired CCR3 usage and partially compromised the efficiency in using CCR5, which could be fully restored by a single escape mutation at the N332 glycan site. Our study demonstrates the link between NAb escape and coreceptor usage alteration in natural HIV-1 infection and indicates that NAb response could drive virus entry tropism evolution *in vivo*.

## Introduction

The highly error-prone nature of HIV-1 reverse transcriptase and rapid virus turn over *in vivo* allow HIV-1 to readily evade host immune response. The ability of HIV-1 to escape neutralizing antibodies (NAbs) *in vivo* is well documented, which renders the contemporaneous viruses relatively resistant to autologous neutralization (Overbaugh and Morris, 2012; Wei et al., 2003). While the impact of the NAb escape mutations on neutralizing sensitivity has been extensively studied, their potential influence on other aspects of HIV-1 phenotype and pathogenesis is much less understood.

The V3 loop of HIV-1 envelope is a major target of autologous NAb response and a key determinant for HIV-1 entry tropism (Cocchi et al., 1996; De Jong et al., 1992; Hwang et al., 1991; Overbaugh and Morris, 2012; Shaik et al., 2019). An early study prior to the identification of the HIV-1 coreceptors showed that the cellular tropism of HIV-1 could change upon escape to V3-specific monoclonal antibodies *in vitro* (McKnight et al., 1995). Because the cellular tropism of HIV-1 is largely governed at the entry level, this observation implied a simultaneous alteration in coreceptor specificity upon immune evasion. Whether this is the case and whether neutralizing response could drive HIV-1 coreceptor usage alteration *in vivo* remain unknown. Indeed, evidence supporting this hypothesis exists. Genetic substitutions at the V3 N332 and N301 glycan sites are typical mechanisms for HIV-1 to evade the V3-glycan specific NAbs (Bonsignori et al., 2017; Krumm et al., 2016; Sok et al., 2016), a class of NAbs with relatively high prevalence in HIV-1 infections (Gray et al., 2011; Landais et al., 2016; Overbaugh and Morris, 2012; Walker et al., 2010). Of note, these glycan sites, especially the N301 glycan are also important in governing HIV-1 coreceptor usage (Ogert et al., 2001; Pollakis et al., 2001). In particular, mutations at the N301 glycan could confer CXCR4 usage when the V3 charge is relatively high (Pollakis et al., 2001). However, the link between NAb escape and coreceptor usage alteration is difficult to demonstrate in the setting of natural HIV-1 infection, largely due to the lack of well characterized cases with both well confirmed escape mutations and clearly defined genetic background the mutations arose.

In the current study, we sought to address the hypothesis that escape mutations selected by autologous NAbs *in vivo* could alter HIV-1 coreceptor usage. To this end, we focused on a previously characterized participant (CH0848) who developed V3-glycan NAb response during natural infection, which selected for viral escape at the N301 and N332 glycan sites (Bonsignori et al., 2017). The well confirmed escape mutations, as well as the availability of viral sequences at acute infection derived by single-genome amplification (SGA) made it possible to precisely determine the impact of escape mutations on coreceptor usage of the cognate T/F virus.

## Material and methods

### Inference of the CH0848 T/F sequence

A total of 1,215 3’-half viral genome sequences previously generated by SGA from participant CH0848 were retrieved from the Los Alamos HIV Sequence Database. The consensus sequence of sequences from the earliest time point (screening) was inferred as the CH0848 T/F sequence as previously described (Keele et al., 2008).

### Pseudoviruses preparation and titration

The CH0848 T/F envelope (*env*) sequence was chemically synthesized (GenScript, USA) and cloned into the pcDNA3.3-TOPO vector (Invitrogen, USA). Immune escape mutations were introduced into the CH0848 T/F *env* by site-directed mutagenesis. All *env* clones were confirmed by sequencing. Pseudovirus stock was prepared as previously described (Song et al., 2016). In brief, 2 μg of each *env* clone was co-transfected with 4 μg of pNL4.3-ΔEnv-vpr+-luc+ into 293T cells in a T25 flask using the FuGENE6 transfection reagent (Promega, USA). The culture supernatants containing the pseudoviruses were harvested at 72 hours post transfection, aliquoted and stored at −80°C until use. The infectious titers (TCID50) of the virus stocks were determined on TZM-bl cells.

### Determination of coreceptor usage

The coreceptor usage of each virus was determined using a panel of NP-2 cell lines expressing CD4 together with different G protein-coupled receptors (CCR5, CXCR4, APJ, CCR3, CCR8, CCR1 and CCR2b) as previously described (Jiang et al., 2011; Song et al., 2016). The parental NP-2 cell line expressing CD4 alone was used as a control to determine the background level of infection. For coreceptor usage assay, NP-2 cell lines expressing different coreceptors were seeded into a 96-well plate one day before infection at a density of 1 × 10^5^ cells per well. On the next day, the cells were infected with approximately 200 TCID_50_ of each pseudovirus (MOI = 0.002). After 6 hours of incubation at 37°C, the infected cells were washed twice with culture medium and cultured at 37°C for three days. At 72 hours post infection, the infected cells were lysed, and the infectivity was determined by measuring the relative luciferase units (RLU) in the cell lysates using the Britelite plus system (PerkinElmer, USA). Viral infectivity was considered positive when the RLU value was at least 5-fold higher than the background RLU value in the parental NP-2 cell line. All experiments were performed in triplicates.

## Results

Participant CH0848 was infected by a single, subtype C T/F virus (Bonsignori et al., 2017) (Fig. 1). V3-glycan specific neutralizing response was elicited in this individual and developed broadly neutralizing activity over the course of infection (Bonsignori et al., 2017). The V3 region of the CH0848 T/F virus contains the conserved N301 and N332 glycan sites (Fig. 2A). Over time, the viruses evaded autologous NAb response by exploring different genetic pathways, including substitutions at the N301 and N332 glycan sites (Bonsignori et al., 2017) (Supplementary Fig. 1). Three escape mutations, N301H, T303I and S334N which disrupted the putative N301 and N332 glycans were selected in the viral quasispecies and were confirmed to escape the recognition by NAb lineages DH270 and DH272 (Bonsignori et al., 2017) (Fig. 2A).

**Fig. 1.**
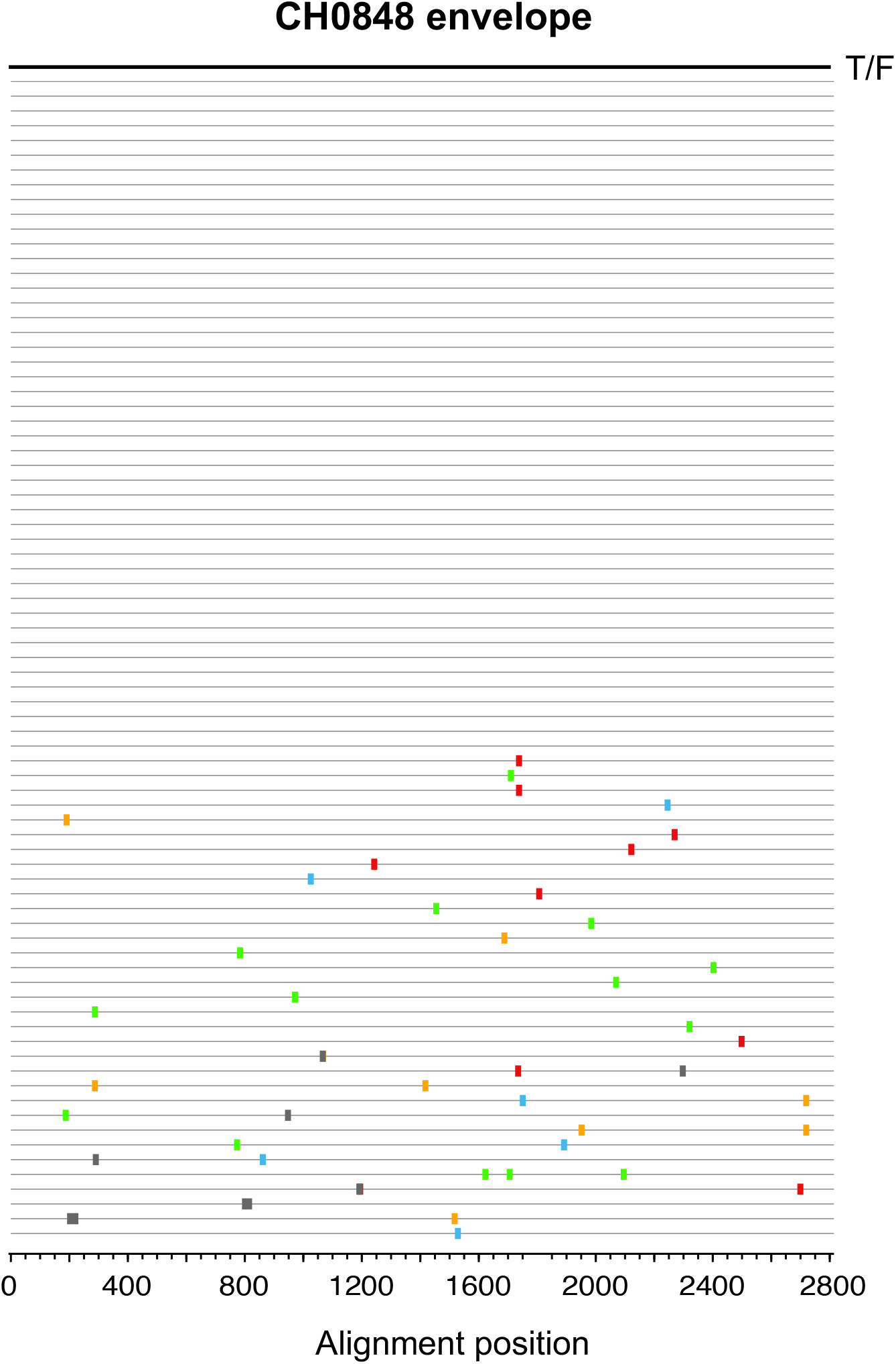
Inference of the CH0848 T/F envelope sequence. Highlighter plot showing sequence alignment of 79 CH0848 envelope sequences from the earliest available time point (screening) generated by single genome amplification (SGA). The consensus sequence (the black line on the top) was inferred as the CH0848 T/F envelope sequence.

**Fig. 2.**
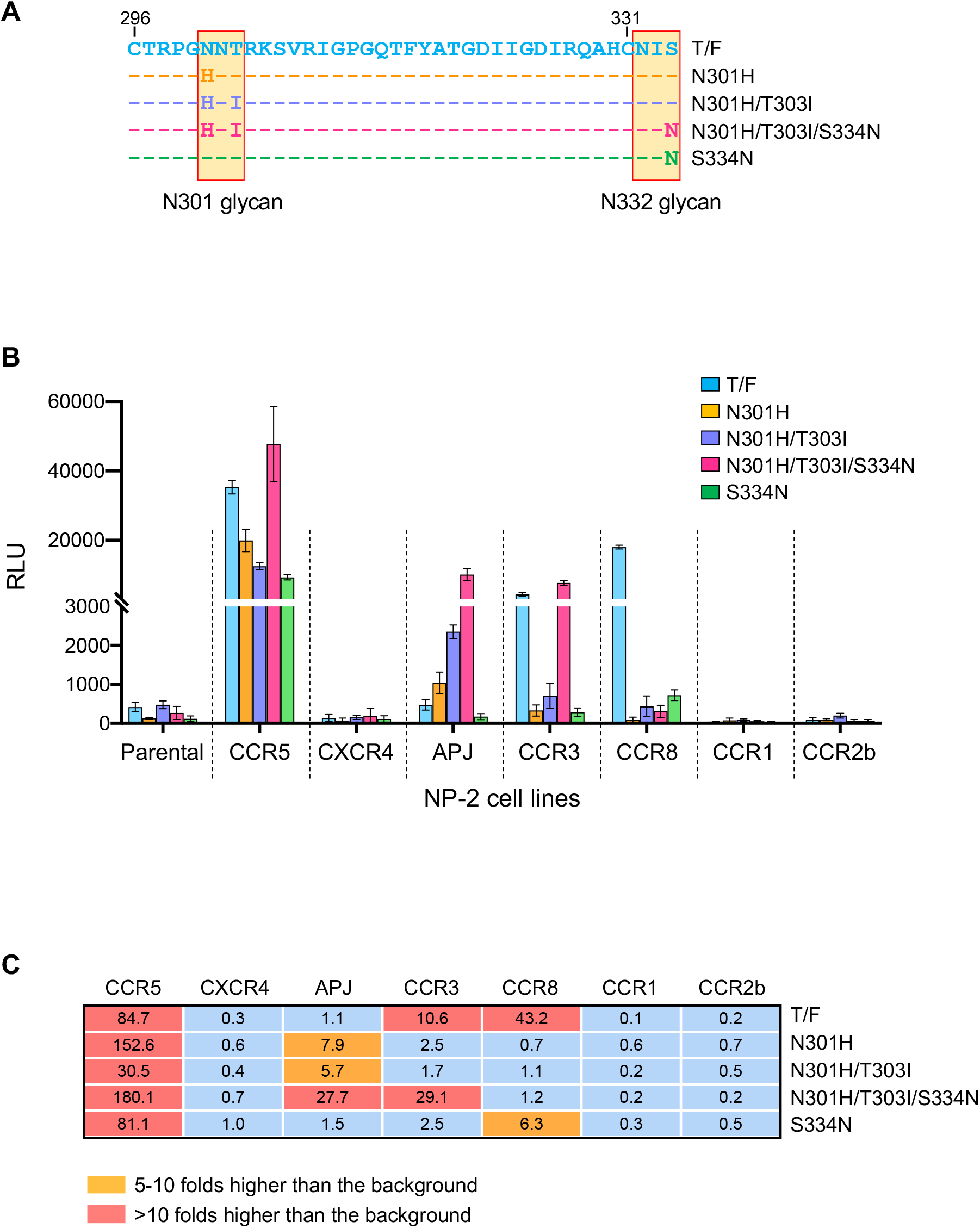
**A.** Amino acid alignment showing the V3 region of the CH0848 T/F virus as well as its escape mutants. The numbers indicate the HXB2 location of the V3 start and end. The N301 and N332 glycan sites were highlighted. **B.** Coreceptor usage determination on NP-2 cell lines expressing CD4 along with a panel of potential HIV-1 coreceptors. Parental cell line expressing CD4 alone was used as a control. All infections were performed in triplicates and the error bars show the standard deviation (SD). **C.** Coreceptor usage repertoire of the CH0848 T/F virus and its escape mutants. The numbers show the fold changes compared to the background RLU value in the parental cell line.

In order to determine the impact of the NAb escape mutations (N301H, T303I and S334N) on the coreceptor usage of the CH0848 T/F virus, we introduced them into the CH0848 T/F envelope individually or in combination (Fig. 2A). Coreceptor tropism assay on NP-2 cell lines expressing CD4 along with a panel of potential HIV-1 coreceptors showed that the CH0848 T/F virus was CCR5-tropic and could use two alternative coreceptors CCR3 and CCR8 for entry (Fig. 2B-C). The efficiency in using CCR8 was relatively high (around 2-fold lower than the infectivity in CCR5 cell line) (Fig. 2B-C). When the impact of different immune escape mutants was assessed, three distinct outcomes were observed. First, mutations at the N301 glycan site (N301H and the N301H/T303I double mutations) completely abrogated the ability to use CCR8 (Fig. 2B-C). The S334N mutation at the N332 glycan site also significantly compromised the CCR8-using efficiency (Fig. 2B-C). Second, the N301H mutation and the N301H/T303I double mutations could confer the ability to use APJ, although the efficiency is relatively low (7.9 and 5.7 folds higher than the background in parental cell line, respectively) (Fig. 2B-C). The S334N mutation itself could not confer APJ usage, but when it was present together with the N301H and T303I mutations, could significantly enhance the efficiency in using APJ (27.7 folds higher than the background) (Fig. 2B-C). This observation indicates that these three mutations may have “synergistic” effect in providing efficient APJ use. Third, the CCR3 usage could be impaired by mutations at the N301 glycan site as well as by the S334N mutation. Interestingly, the level of CCR3-mediated entry was fully restored when the three mutations were present together (Fig. 2B-C).

Previous study showed that mutation disrupting the N301 glycan could abrogate CCR5 usage of a dual-tropic HIV-1 isolate (Ogert et al., 2001). In the genetic background of the CH0848 T/F virus, all escape mutants maintained CCR5 usage. The N301H/T303I double mutations could partially compromise the efficiency in using CCR5, but the S334N mutation could restore the CCR5 usage to the wild-type level (Fig. 2B-C). This observation indicates that in addition to immune escape, the S334N mutation may have a compensatory effect to maintain efficient CCR5 usage when the N301 glycan was disrupted. This could explain why mutations at the N301 glycan site tended to form linkage with the S334N mutation in the viral quasispecies in CH0848 (Supplementary Fig. 1). Together, these observations demonstrate that immune escape mutations at the N301 and N332 glycan sites selected by V3-glycan NAbs *in vivo* can alter the coreceptor usage repertoire of the T/F virus.

## Discussion

Our study demonstrates the link between NAb escape and coreceptor usage alteration in HIV-1 infection. As immune escape to NAb response inevitably occurs during natural HIV-1 infection, the biological and pathogenesis consequence of the simultaneous change in coreceptor usage require further investigation with larger number of samples from different HIV-1 subtypes.

Previous studies showed that mutations occurring at the N301 glycan site could abolish the CCR5 usage of dual-tropic HIV-1 and confer CXCR4 usage on R5-tropic HIV-1 (Ogert et al., 2001; Pollakis et al., 2001). Our previous study on CRF01_AE infections showed that all X4-using viruses isolated from HIV-1 infected people contained mutations at the N301 glycan site (Song et al., 2019). In the genetic background of the CH0848 T/F virus, NAb escape mutations at the N301 glycan site could abrogate CCR8 usage while confer efficient APJ usage when present together with the S334N mutation. These observations together indicate the important role of the N301 glycan site in governing HIV-1 coreceptor specificity and suggest that the exact impact of the mutations could depend on the genetic backgrounds and HIV-1 subtypes. It is possible that the process of “coreceptor switch” could be one particular outcome among different patterns of coreceptor usage alteration driven by NAb response. Future studies by investigating the co-evolution between the virus and NAbs in HIV-1 infected individuals with coreceptor switch will be important to address this possibility.

Although the significance of alternative coreceptor usage in HIV-1 pathogenesis and transmission remain to be determined, the biological consequence upon the acquisition of APJ use by the escape mutants deserves further investigation. The wide expression of APJ in brain has raised the possibility that it could contribute to the neuropathogenesis of HIV-1 (Cayabyab et al., 2000; Edinger et al., 1998). Our previous study has identified a T/F HIV-1 which could not use CCR5 efficiently but could use GPR15 and APJ with comparably high efficiency (Jiang et al., 2011), indicating a potential role of APJ in HIV-1 transmission. Although previous studies suggested that the V1V2 region may play a major role in determining APJ use (Hoffman et al., 1998), the current study indicates that the V3 glycan sites could also be important. The observation that mutations at the N301 and N332 glycan sites have a “synergistic” effect in providing efficient APJ usage indicates that the loss of V3 glycosylation might favor APJ recognition. It is also possible that the substitutions in the amino acid backbone, rather than the loss of glycosylation itself contribute to APJ usage.

Beyond the HIV field, our study may have implications for understanding the molecular mechanisms driving virus receptor specificity alteration in natural host. The fast-evolving nature and high genetic plasticity confer receptor usage flexibility on many viruses, from bacteriophage to human viruses (Baranowski et al., 2001; Baranowski et al., 2000; Berger et al., 1999; Liu et al., 2002). Alteration in receptor specificity could be closely related with disease progression, as the process of HIV-1 coreceptor switch (Berger et al., 1999; Regoes and Bonhoeffer, 2005; Schuitemaker et al., 2011; Verhofstede et al., 2012), and could contribute to cross-species viral transmission, as exemplified by coronavirus and influenza virus (Parrish et al., 2008; Peck et al., 2015; Shi et al., 2014). Although in experimental settings, minimal genetic changes could alter virus receptor specificity, the “*bona fide*” process happening in natural host is difficult to capture and the underlying molecular mechanisms remain a major knowledge gap. Given that the overlap between neutralizing epitopes and receptor binding regions is widely observed for viruses from different families (Fields et al., 2013; Howley and Knipe, 2021), it is possible that like the V3 glycan sites in the HIV-1 genome, “dual determinants” for both receptor specificity and NAb susceptibility exist in other viruses as well. From an evolutionary perspective, a simultaneous shift in receptor usage upon immune escape could provide the virus an alternative route to maintain its entry ability while evading the host immune defense (here, we term this hypothesis the “escape by shifting” concept). Further studies by exploring the “escape by shifting” scenario in different viral infections could provide novel insights into viral pathogenesis, transmissibility, as well as zoonotic potential.

## Author contributions

Manukumar Honnayakanahalli Marichannegowda: Investigation, Formal analysis, Data Curation. Hongshuo Song: Conceptualization; Investigation; Supervision; Writing – Original Draft.

## Declaration of competing interest

The authors declare that they have no conflict of interest.

## Acknowledgements

This work was support by the Institute of Human Virology, University of Maryland School of Medicine.

## Figure legends

**Supplementary Fig. 1.** Longitudinal amino acid sequences showing the evolution of the CH0848 V3 region. All sequences were downloaded from the Los Alamos HIV Sequence Database. The N301 and N332 glycan sites are highlighted in green.

## Notes

### Competing Interest Statement

The authors have declared no competing interest.

